# A Rapid and Scalable Subcutaneously Administered Murine Thymus Micro-organoid for Generating Functional T cells

**DOI:** 10.1101/2025.09.05.671099

**Authors:** Simon Gebremeskel, Inah Bianca Embile, Nikolay Bazhanov, Chuo Fang, Ashley Heard, Jinsam Chang, Christen Boyer, Midori Taruishi, Phillip Baker, Kelsey Howard Taylor, Pete O’Heeron, Hamid Khoja

## Abstract

Thymic function can decline due to age-related involution, congenital disorders, acute infections, or chemo/radiation therapy. Decline in thymic function leads to decreased T cell production and weakened immunity. To address these thymic insufficiencies, we aimed to develop a transplantable and scalable micro-organoid system utilizing fibroblasts and thymic cells. We have developed a reliable and rapid method to generate thymic micro-organoids using selectively screened fibroblasts and murine thymic cells. The thymic micro-organoids are cryo-preservable, injectable, and give rise to T cells both *in vitro* and *in vivo*. Thymic organoids expressed key genes required to sustain T cell development and maturation: *ccl25, dll-1, dll-4, foxn-1, il-7, scf*. When injected into T cell-deficient *Prkdc^scid^* mice, the organoids gave rise to functional αβ, ψ8, natural killer T (NKT) cells, and FoxP3^+^ regulatory T cells. Organoid-derived T cells expressed a diverse T cell receptor (TCR) repertoire i*n vivo* and responded to stimulation with anti-CD3/28, Concanavalin-A, or Phytohemagglutinin. Thymic organoids derived from pmel-1 thymocytes gave rise to Vβ13^+^ T cells that delayed the growth of B16 melanoma and enhanced activation of T and NK cells. This approach presents a valuable tool for mechanistic studies and addressing current therapeutic gaps in diseases associated with thymic decline and insufficiencies.

## Introduction

The thymus is a lymphoid organ where precursors of T cells undergo differentiation, selection, and proliferation. This process gives rise to diverse, functional, and self-tolerant T-cell populations. Age-related thymic involution is a natural process that leads to a gradual decrease in thymic size, function, and T cell production capabilities. In humans, age-related thymic involution begins after birth, and the size of the thymus peaks during puberty, before gradually decreasing. Thymic involution results in a progressive decline in the production of naive T cells, a reduction in T cell receptor (TCR) diversity, and impaired immune responses. In addition to this natural decline in thymic function, the proper functioning of the thymus can also be impaired by infections, exposure to chemotherapy or radiation therapy, fluctuations in hormone levels, and many other idiopathic causes (1, 2). In some patients, complete athymia may result from congenital diseases such as FOXN1 mutation, DiGeorge Syndrome, diabetic embryopathy, and idiopathic causes (3, 4).

Decreased thymus function or complete thymectomy results in susceptibility to cancer, autoimmune diseases, infections, and poor immunity following vaccination (1, 2, 5). In a retrospective study with 1420 thymectomy patients and 6021 control patients, thymectomy patients had a higher risk of mortality from all causes and a higher incidence of cancer (5). Highlighting the need for additional therapeutic interventions. Currently, complete athymia is treated with an allogeneic thymus transplant or hematopoietic stem cell transplant (3, 4). Despite advances in our understanding of T cell biology, model systems to interrogate certain aspects of T cell development or the mechanisms underlying T cell insufficiencies remain understudied. Experimental model systems to address these gaps in knowledge are needed and may help to facilitate the development of T cell therapeutics.

The early thymic progenitors (ETPs) originate from the fetal liver or post-natal bone marrow and enter the thymic cortex as double-negative thymocytes (CD4^-^ CD8^-^, DN), via the high endothelial venules in the cortico-medullary junction. In the outer cortex, the ETPs rapidly proliferate, express a TCR, and undergo positive selection. The interaction between the TCR and MHC complexes on thymic epithelial cells (TECs) triggers positive selection, leading to differentiation into single positive CD4^+^ or CD8^+^ thymocytes with functionally diverse TCRs. Positively selected thymocytes then migrate from the cortex to the medulla, where they are exposed to tissue-restricted antigens and undergo negative selection of self-reactive T cells and/or the differentiation into regulatory T cells (Tregs). These processes are guided in part by a rich assortment of thymic stromal cells. The thymic stroma consists of non-hematopoietic cells such as thymic mesenchymal cells, TECs, and cells of hematopoietic origin, including dendritic cells, B cells, and macrophages (6). The thymic stroma provides a foundation on which T cell precursors can nest, mature, and eventually enter the periphery.

*In vitro* studies of thymus biology and T cell development are challenging, in part because two-dimensional cultures fail to maintain their functionality (7, 8) and TECs are difficult to propagate in culture. There have been recent advances to recapitulate certain aspects of thymus biology *in vitro*, including OP9/MS5-DLL1/DLL4 feeder cell systems (9, 10), thymus re-aggregate cultures, and fetal thymus organ cultures to model thymus organogenesis (11–16). Many of these approaches rely on Matrigel or similar bioproducts, require extended duration cultures, and face limitations in large-scale production. We sought to address some of these existing challenges.

Fibroblasts play an important role in maintaining the structural architecture of organs within our body. Using Fibroblasts, we sought to develop a reproducible, timely, cost-effective, matrix-free, and scalable approach to developing thymic organoids in culture. By mixing dissociated thymic cells with fibroblasts, we were able to develop an approach to rapidly generate murine micro-organoids that can give rise to T cells when transplanted into mice. The size of the micro-organoids can be modulated to different sizes and can be cryo-preserved for long-term storage. When transplanted into mice, they gave rise to conventional CD4^+^ and CD8^+^ T cells, FOXP3^+^ Tregs, γδ T cells, andg NKT cells. Akin to the native thymus, the micro-organoids gave rise to T cells with diverse TCR and responded to stimulation, a key requirement for T cell-mediated immunity. Furthermore, Pmel-1 thymocytes cultured in micro-organoids gave rise to Vβ13^+^ T cells that delayed B16 melanoma growth in mice and resulted in activated T and NK cells in the draining lymph nodes.

## Materials and Methods

### Mice

BALB/c (#000651), BALB/c scid (#001803), C57/BL6 (#000664), pmel-1 (#005023) mice (6-8 weeks) were purchased from Jackson Laboratories and maintained at Stillmeadow Animal Care facility (Sugar Land, TX). Mice were housed in temperature-controlled rooms with a 12-hour light/dark cycle and had access to food and water ad libitum. All experimental procedures were reviewed and approved by the Stillmeadow Institutional Animal Care and Use Committee.

### Cell culture

Mouse dermal fibroblasts (MDF) were sourced from ATCC and cultured in DMEM-F12 (Gibco) supplemented with 10% FBS (Avantor) and 1% Pen-Strep (Gibco) (Complete media). B16-F10 mouse melanoma cells were purchased from ATCC and cultured in DMEM (Corning) supplemented with 10% FBS and 1% Pen-Strep. Cells were cultured at 37°C, 5% CO_2_.

### Thymus processing

Thymus tissue was harvested from 6–8-week-old mice and digested in 200 μg/ml Liberase (Roche) for 30-45 min at 37°C. Cells were then passed through a 70 μm cell strainer, washed, and counted prior to use in subsequent experiments.

### Thymus organoid formation

Fibroblasts were cultured to 80% confluency and harvested using 0.25% trypsin for 5 minutes. Cells were washed and resuspended in complete media. We combined fibroblasts with dissociated thymus tissue at varying ratios (1:6, 2:6, and 3:6 fibroblasts to thymic cells) and plated them in ultra-low attachment plates for 72 hours. After 72 hours, the organoids were used for *in vivo* and *in vitro* characterization. Each organoid contained approximately 6000 thymic cells and had an average diameter of 175 µm. Supernatants were obtained from thymus organoid cultures for cytokine evaluation. Stem cell factor (SCF) levels were quantified using an ELISA in accordance with the manufacturer’s instructions (Invitrogen, Cat# EMKITL).

### Fluorescent dye staining

In some experiments, MDFs and thymocytes were labeled with Calcein Am dye (ThermoFisher) and Eflour 670 dye (eBiosciences) according to the manufacturer’s recommendations. Briefly, MDFs and thymocytes were washed twice in PBS. The MDFs were then resuspended in 1µM Calcein AM, whereas thymocytes were resuspended in 5µM eFlour 670 dye. Cells were incubated at 37°C for 10 minutes, and staining was stopped by adding an equal volume of FBS. Cells were then washed twice in media containing 10% FBS and used for organoid cultures.

### Flow cytometry

The following antibodies were purchased from BD Bioscience, eBioscience, or Biolegend (clone names are in brackets): purified CD16/32 (97), 7-AAD viability stain; FITC-labeled CD4 (RM4-5); PE-Labeled CD8α (53-6.7), FOXP3 (FJK-16s), TCR-γδ (GL3); APC-labeled CD3 (145-2C11), APC-eFlour780-labeled TCR-β (H57-597), NK1.1 (PK136); Brilliant Violet 421-labeled CD25 (PC61); Brilliant Violet 510-CD45.2 (104). Cells were pre-incubated with CD16/32 Fc blocking cocktail and stained with surface staining antibody panel and incubated at 4°C for 20 minutes, washed, and fixed in 2% paraformaldehyde. For intracellular staining, cells were stained using the BD Bioscience FOXP3 staining kit according to the manufacturer’s instructions. Analysis was performed using a Cytoflex flow cytometer, and data analysis was performed using FlowJo software.

### Viability assay

Thymic organoids were harvested, rinsed twice in PBS, and dissociated in Accumax (ThermoFisher) for 30-50 minutes on a plate shaker at 1400rpm. The dissociated cells were neutralized in complete medium and rinsed with PBS. Cells were stained with 7-AAD, and data were acquired on a Cytoflex flow cytometer.

### In vivo models

To examine whether thymic organoids could give rise to mature T cells *in vivo*, we injected 5000 organoids (containing 3×10^7^ thymocytes) subcutaneously into immune-deficient SCID mice. On days 25 and 42, blood was obtained via submandibular venipuncture: 70 µl of blood was collected into EDTA-containing microtubes. We analyzed the frequency of CD45^+^7AAD^-^ TCRβ^+^ T cells, CD4^+^ T cells, and CD8^+^ T cells in peripheral blood. At the experimental endpoint, we evaluated the frequency of splenic αβ T cells, CD4^+^ T cells, CD8^+^ T cells, FOXP3^+^ regulatory T cells, and γδ T cells. We further evaluated the TCR diversity of CD3^+^ T cells using the BD Pharmingen mouse Vβ screening kit.

To examine whether CD3 T cells would reconstitute all organs, we harvested tissue from the spleen, liver, lung, kidney, and heart for transcript analysis.

### Immunohistochemistry

Samples were formalin-fixed and embedded in paraffin blocks. The embedded tissues were sectioned into 4-5 µm cryostat sections and mounted onto slides for immunohistochemical analysis. Deparaffinization was performed by immersing slides in Histo-Clear for 5 minutes, repeated twice. Tissue sections were rehydrated in graded ethanol concentrations: 100%, 90%, 80% and 70%, followed by distilled water with each step lasting for 3 minutes. Antigen retrieval was performed in 1% Citrate (Vimentin staining) or EDTA (CD3 staining) at 95°C for 30 minutes. Slides were rinsed with PBS-Tween for 2 minutes each and exposed to 3% hydrogen peroxide for 10 minutes to inactivate endogenous peroxidase activity. Samples were blocked in PBS containing 1% BSA and 0.3% Triton X-100 for one hour at room temperature, followed by overnight staining with either rabbit anti-CD3 (Abcam 135372) or rabbit anti-Vimentin (Abcam ab92547) antibodies prepared at a 1:200 dilution in blocking buffer. The samples were rinsed three times in PBS with 5-minute incubations and stained with secondary antibody (1:1000) for 1 hour at room temperature. Sections were washed in PBS, incubated in ABC solution (Vector Laboratories) for 30 minutes, and washed again. DAB substrate (Vector Laboratories) was applied for 2–3 minutes until the desired staining developed, then stopped with distilled water. Slides were counterstained with Mayer’s hematoxylin (Abcam) for 5-10 seconds, rinsed, dehydrated through graded ethanol concentrations and Histo-Clear, and mounted with coverslips using Toluene mounting medium. Brightfield images were acquired using the Echo Revolve microscope (Discover Echo).

### T cell proliferation

To examine whether T cells derived from thymus organoids were functional, we isolated splenocytes from SCID mice reconstituted with thymus organoids. Splenocytes were labelled with CFSE dye according to the manufacturer’s recommendations and stimulated in culture with T cell activator CD3/28 dynabeads (Gibco), Concanavalin A (Con A) (Invitrogen), or phytohaemagglutinin (PHA) (Invitrogen). We examined T cell proliferation via CFSE dilution after 96 hours of stimulation. IFN-γ production was quantified via ELISA (Invitrogen 88731422) in accordance with the manufacturer’s instructions.

### Tumor model

Next, we examined whether organoid-derived T cells could protect mice from tumors. To test this, we isolated thymic tissue from pmel-1 mice and formed organoids in culture. Prior to transplanting organoids into mice, we preconditioned mice with 150 mg/kg (i.p) cyclophosphamide for three consecutive days. One week following transplantation, we inoculated mice with B16 melanoma and monitored tumor growth for up to 2 weeks. Tumor volume was calculated as 0.5*a*b^2^ (where a is the length, and b is the width of the tumor) (17). On day 14, tumors and lymph nodes were harvested and dissociated into single cells using mechanical dispersion through 70 μm filters. Leukocytes were enriched using a 30% Ficoll gradient centrifugation at 500g for 20 minutes. After collecting the leukocyte-containing buffy coat, the remaining red blood cells were lysed using RBC lysis buffer (eBiosciences) and washed with PBS containing 2% FBS. Cells were phenotyped using a Cytoflex flow cytometer (Beckman Coulter) and counted using the NC3000 cell counter (Chemometec).

### RNA isolation

Total RNA was isolated using the RNeasy kit (Qiagen), and cDNA was prepared from 500 ng of total RNA using the high-capacity cDNA reverse transcription kit (Applied Biosystem). Relative Quantitative PCR was performed in replicate using 1 µL of cDNA and Taqman primers. PCR was run with 10 10-minute hold at 95°C before 40 cycles of 15 seconds at 95°C followed by 45s at 55°C. Data were collected on a Quant Studio 6 (Applied Biosystem) and relative expressions to house-keeping control gene (β-actin) were derived using the 2−ΔCT relative quantification technique. Validated high-stringency TaqMan primers were used for: *β-actin* (Mm02619580_g1)*, cd3*e (Mm01179194_m1), *ccl25* (Mm00436443_m1)*, delta like ligand 1 (dll1)* (Mm01279269_m1)*, delta like ligand 4 (dll4)* (Mm01338018_m1)*, forkhead box protein n1 (foxn1)*(Mm00433948_m1)*, stem cell factor (scf)* (Mm00442972_m1)*, interleukin-7 (il-7)* (Mm01295803_m1).

### Statistical analysis

All data were acquired from three independent experiments and are expressed as mean ± SEM. A two-tailed Mann-Whitney *U* test was used to compare the two groups. Comparisons between more than two groups were made via Kruskal–Walli’s analysis with Dunn post-test. Tumor volumes were evaluated using a mixed-effects model with a Fisher’s least significant difference post hoc test. Statistical significance was set at *P* < 0.05, and statistical computations were performed using GraphPad Prism 10.02. Image J software (NIH) and Biorender software were used to prepare colored or schematic images.

## Results

### Characterizing thymic micro-organoids

Reaggregate thymus cultures and miniature thymic organoids have been previously described (11–13, 15, 18). These approaches required long-duration cultures, the use of bioproducts such as Matrigel, and were difficult to scale up. Given the important role fibroblasts play in tissue architecture, we hypothesized that fibroblasts may help address these challenges and allow for the scale-up production of thymic organoid cultures. To test this approach, we dissociated thymic tissue into single cells and co-cultured the dissociated thymus cells with fibroblasts in ultra-low attachment plates (Figure 1A). Dissociated thymic cells on their own did not form micro-organoids after 3 days (Figure 1B). However, thymic cells combined with fibroblasts were able to reproducibly generate thymus micro-organoids (Figure 1B). This suggests that fibroblasts play a critical role in the structural formation of the thymic organoids. The size of the organoids can be modified by altering the number of fibroblasts added to each well while maintaining a constant 6000 thymic cells in each organoid (Figure 1C). Larger organoids result in increased cell death at the core (19, 20), and may disaggregate when injected through a needle. Therefore, selecting an optimal size range for the organoids was critical. To examine the distribution of thymic cells and fibroblasts, we pre-labelled thymic cells with eFluor 670 membrane dye and fibroblasts with Calcein AM dye. The labeled cells were then combined to form thymus organoids and imaged after 3 days in culture. Fibroblasts and thymus cells were evenly distributed throughout the organoids, with no discernible compartmentalization (Supplementary Figure 1).

**Figure 1.**
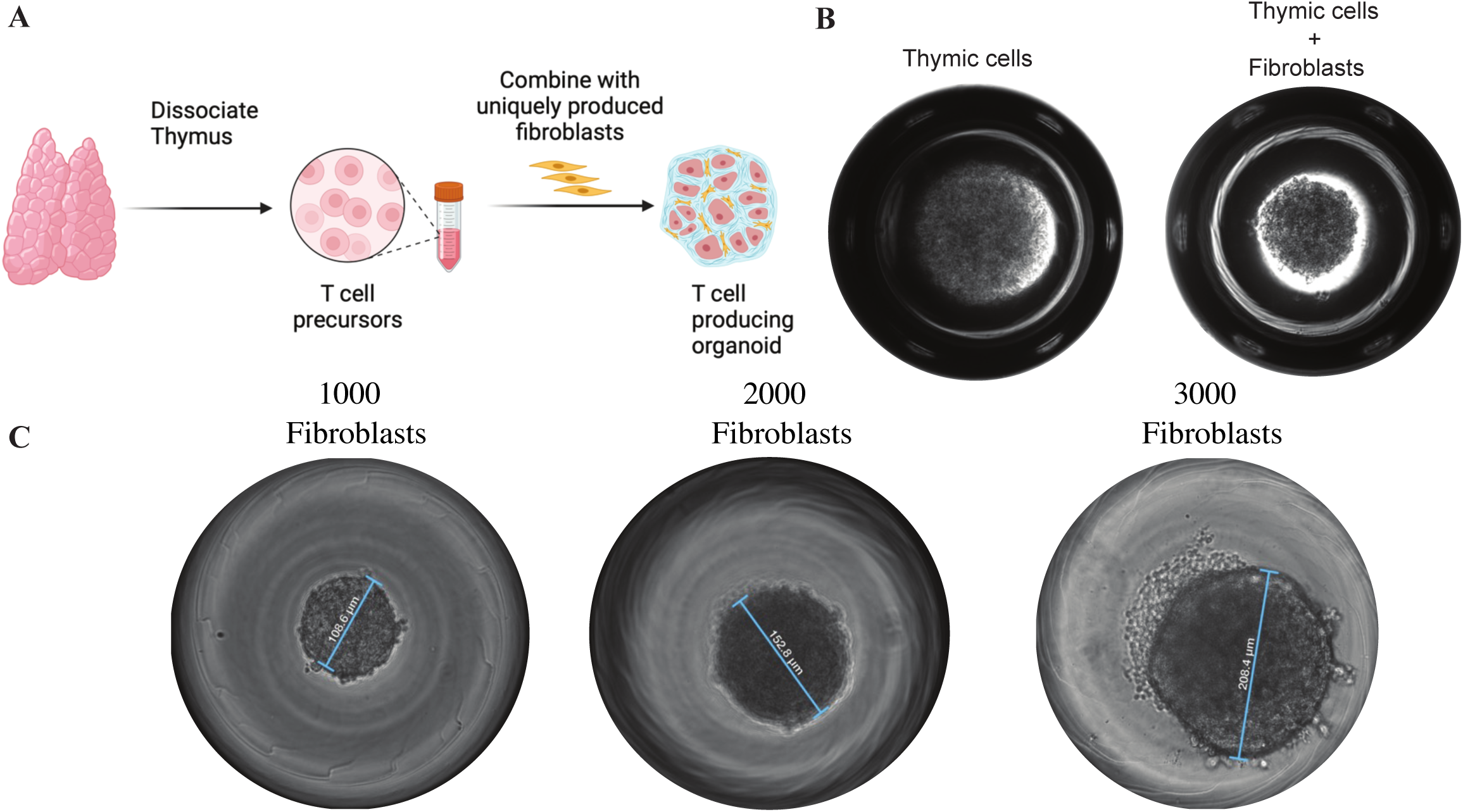
Thymus micro-organoid culture. A) A schematic depicting the formation of the thymus micro-organoids. Thymus tissue is dissociated into single cells, combined with fibroblasts, and seeded into a low-binding plate. B) Brightfield image of thymus micro-organoids taken 3 days post-seeding. C) Fibroblast concentration corresponds with the size of the micro-organoids. Increasing concentration of fibroblast (1×10^3^, 2×10^3^, 3×10^3^) while keeping constant thymocyte numbers (6×10^3^).

We next examined whether the thymus organoids could give rise to differentiated T cells. After 3 weeks in culture, we were able to detect double-positive T cells (Figure 2A). However, further culturing the organoids did not result in differentiation into single positive T cells. It is worth noting that no exogenous cytokines were added to the culture. We next examined whether the organoids expressed transcripts of key cytokine and growth factors required for T cell development. We isolated total RNA from the thymic organoids grown in vitro, and we were able to detect transcripts that are critical for T cell development (*ccl25, dll1, dll4, foxn1, scf, il-7*) (Figure 2B), suggesting that the organoid could sustain T cell development when transplanted into mice. We confirmed the protein expression of SCF in the culture supernatant. Interestingly, fibroblasts appear to be the primary source of SCF in the organoids (Figure 2C). This suggests that, in addition to the aforementioned contribution of fibroblasts to the structural integrity of the thymic organoids, fibroblasts also produce growth factors that enhance the survival of T cell precursors. Organoid cultures typically take several weeks and require constant monitoring and maintenance (11–16). We therefore examined whether the culture period required to obtain a viable product could be shortened. We found that within 72 hours of culture, the thymic micro-organoids can be harvested and transplanted into mice (Figure 1B).

**Figure 2.**
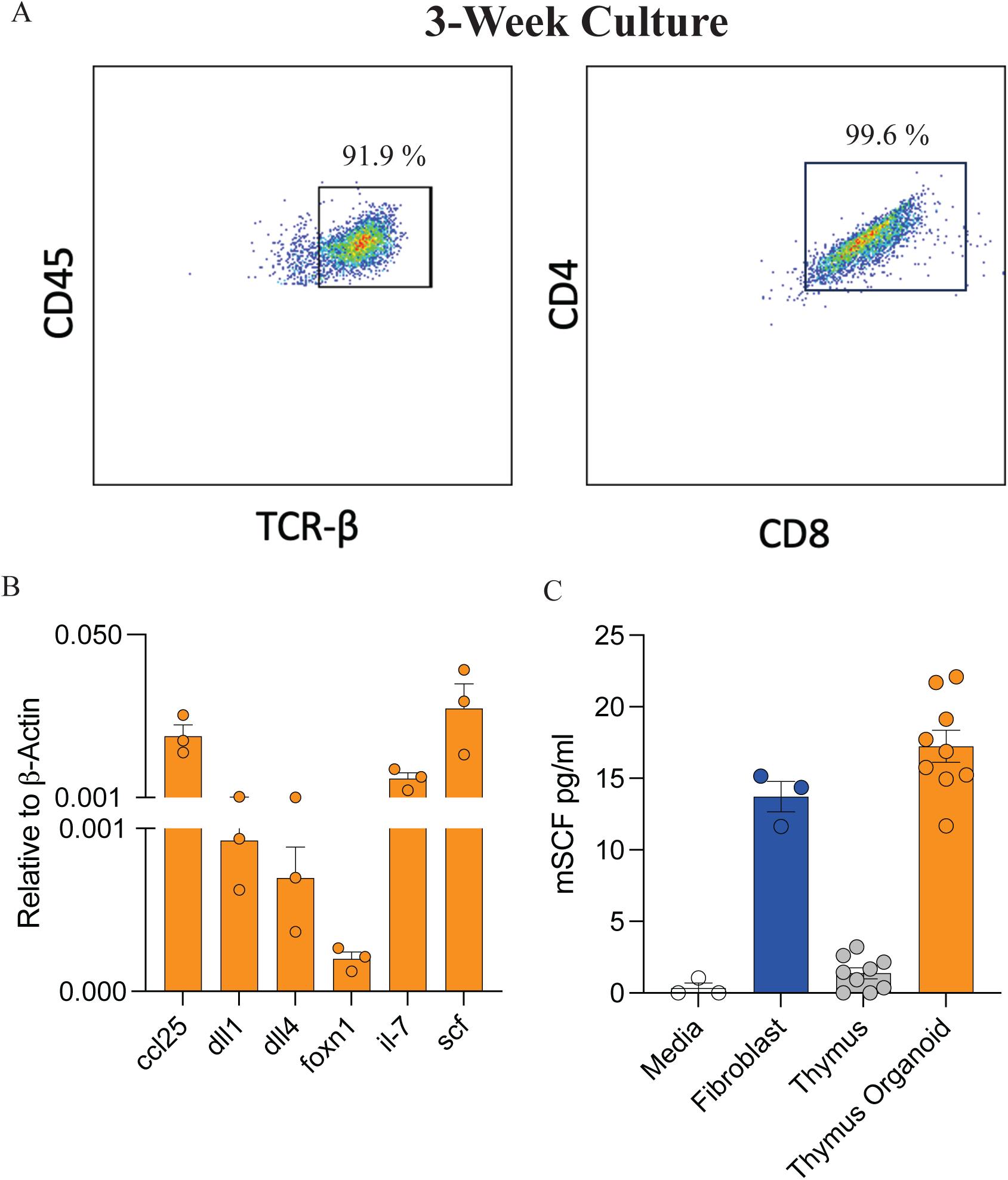
Thymus micro-organoids maintained for more than 4 weeks in culture without addition of exogenous cytokines or growth factors. A) After 3 weeks in culture, the majority of thymocytes are DP (CD45^+^ TCR^+^ CD4^+^ CD8^+^). Numbers indicate percentage of double-positive cells. B) Thymus organoids express key genes required for T cell development and function: Cytokines, growth factors and notch ligand mRNA expression in thymus organoid after 3 weeks of culture. Relative gene expression was analyzed by comparing to the validated housekeeping gene δ-actin (2−ΔCT) (n=3).

### Organoid transplantation

Current thymus transplant strategies require invasive surgery in immunocompromised infants (3, 4). We wanted to examine whether our organoids could be delivered without surgery, avoiding some of the complications and costs associated with surgery. Given that we can modulate the size of the organoids, we optimized the cultures for micro-organoids that were less than 200 μm in diameter (Figure 1C). This enabled the micro-organoids to be easily loaded into syringes and injected with a 27-gauge needle without disrupting the integrity of the micro-organoids. We transplanted thymic organoids subcutaneously into immune-deficient SCID mice. Circulating T cells were detected at low levels as early as two weeks post-transplant (Supplementary Figure 2), and the frequency was higher at week 4 and 6 (Figure 3A). To determine whether the reconstituted T cells resembled wild-type mice, we compared the frequency of CD4^+^ and CD8^+^ T cells in the reconstituted mice to those in wild-type mice. There was a slight trending elevation in the frequency of CD4^+^ T cells and a trending decrease in CD8^+^ T cells when compared to wild-type mice, but this did not reach statistical significance (Figure 3C). Further detailed monitoring was precluded, as some of the mice began to show signs of colitis on day 52, leading to an early endpoint in the study. The development of colitis has been well established in T cell transfer models, where splenic T cells are adoptively transferred into immunodeficient mice (21). While this was not a stated objective of our study, it points to the functionality of T cells derived from the thymus organoid.

**Figure 3.**
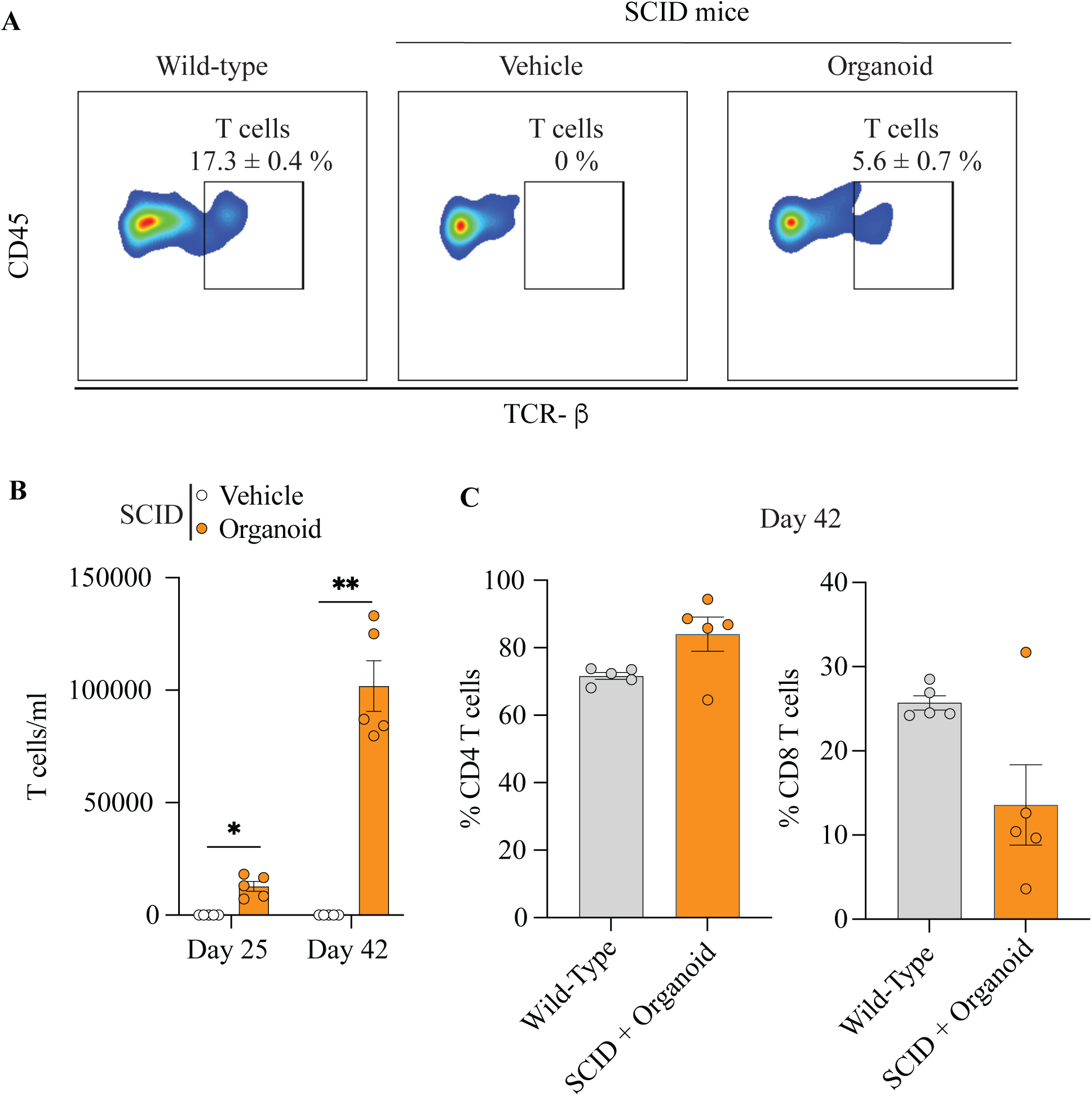
Organoids reconstituted T cells in immunodeficient SCID mice. A) Representative flow cytometry plots of circulating T cells (TCR^+^ CD45^+^) in wild-type mice, naive SCID mice, and SCID mice transplanted with thymus organoids. Numbers indicate percentage frequency. B) Number of circulating T cells per volume of blood on day 25 and 42 following thymus organoid transplantation in SCID mice (n=5). C) Frequency of CD4^+^ T cells and CD8^+^ T cells in wild-type mice and SCID mice transplanted with thymus organoids (n=5).

### Cryopreservation

Current approaches to thymic organ transplant require the maintenance of fresh organ culture for up to 24 days (3, 4, 22). Thymic organoids in this study were cryopreserved in freezing media containing 5% DMSO (1:1 mixture of cryostor 10 and plasmalyte A) and maintained high viability upon thawing (Figure 4A). When Organoids were transplanted into mice, they gave rise to viable T cells, indicating that frozen thymus organoids retained their capacity to generate T cells. There was no difference in the T cell reconstitution levels between mice receiving fresh or cryopreserved organoids (Figure 4B). This observation marks a significant advancement in the potential for organoid transplantation, as current clinical approaches often rely on fresh cultured organoids.

**Figure 4.**
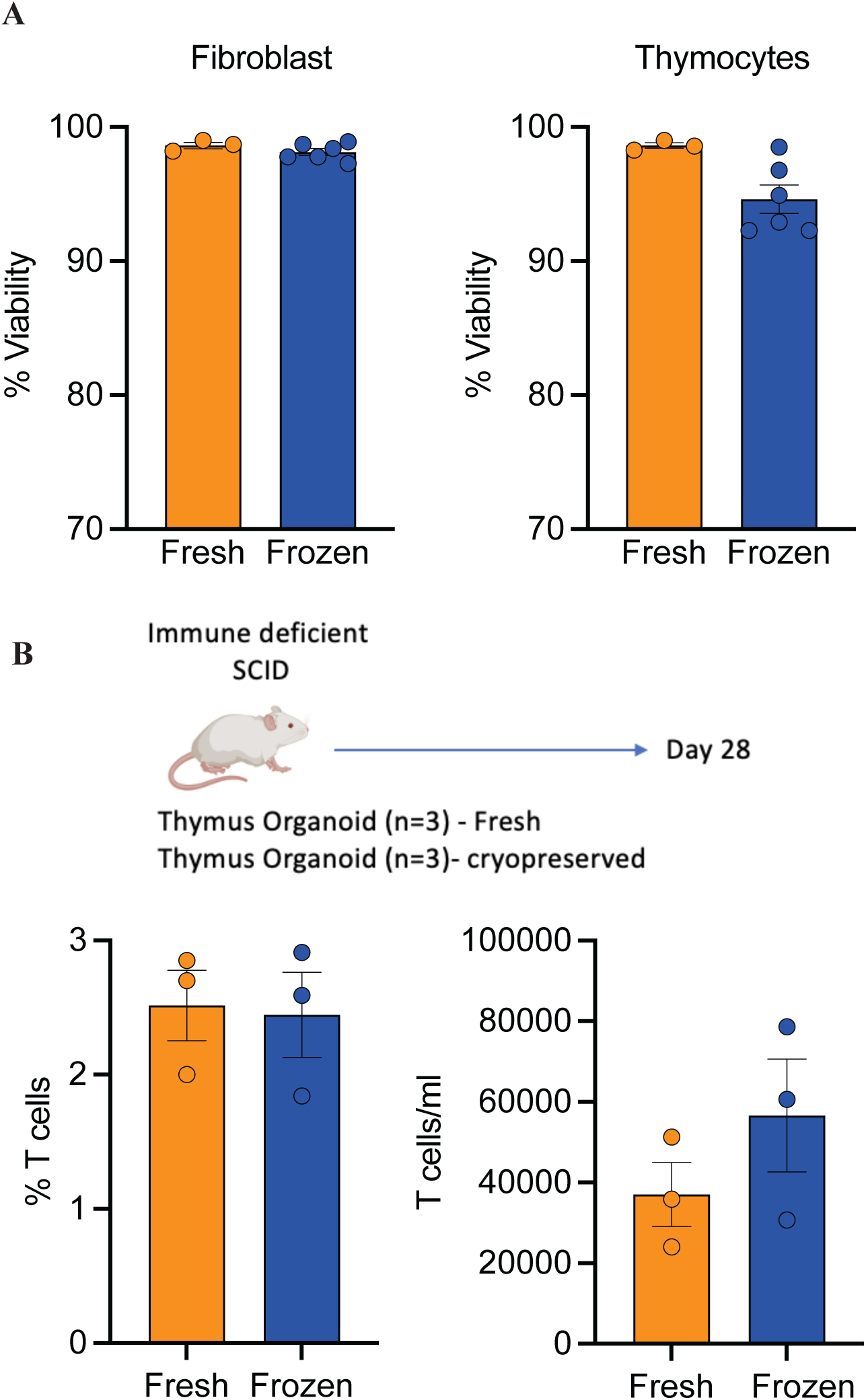
Cryopreserved organoids were viable and maintained T cell-producing capacity. A) Cryopreserved thymus organoids maintained high viability upon thawing (n=3). Thymus organoids were cryopreserved at -80 °C in freezing medium containing 5% DMSO. Cells were thawed, dissociated into single cells, and stained with 7AAD to determine viability using a flow cytometer. B) Frequency and number of circulating T cells in SCID mice transplanted with fresh or cryopreserved organoids (n=3). Organoids were transplanted subcutaneously into SCID mice, and 28 days following transplantation, T cells in circulation were monitored via submandibular venipuncture.

### TCR diversity

T cells activation occurs primarily through recognition of antigens via the TCR. To recognize a broad array of potential danger signals, mammals have evolved to express a highly diverse TCR repertoire. The thymus plays a critical role in ensuring T cell precursors undergo successful TCR rearrangement, resulting in T cells that recognize foreign antigens but do not attack cells expressing self-antigens. To examine TCR diversity, we screened for common TCR Vβ profiles using flow cytometry. Thymic organoids gave rise to T cells expressing a diverse TCR Vβ profile (Figure 5A), implying there was no selective pressure or bias towards a subset of TCR Vβ chains. This suggests that the T cells generated will likely be able to recognize a broad array of antigens.

**Figure 5.**
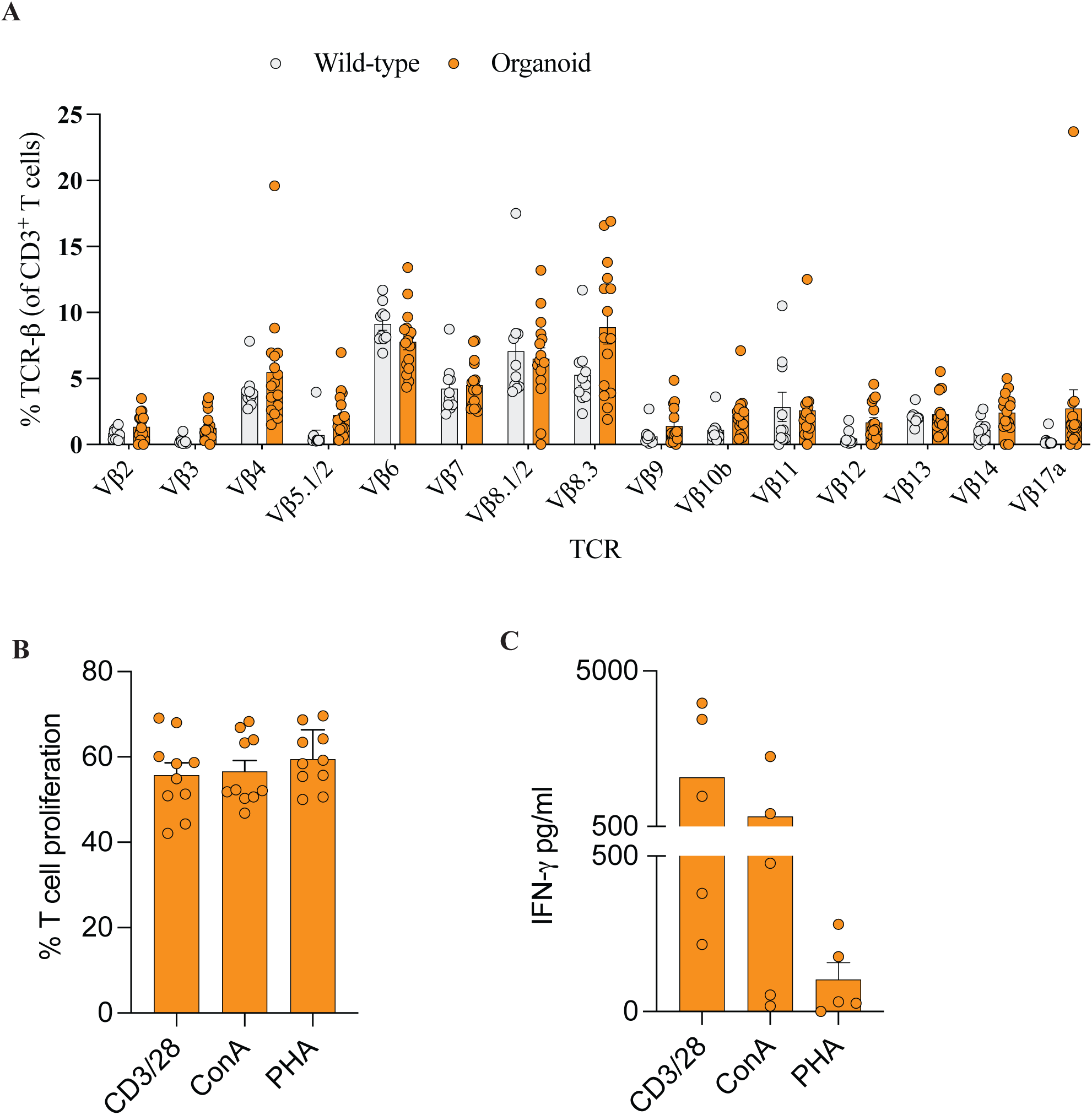
Transplanted organoids gave rise to diverse T cells that respond to stimulation. A) Vβ profile of splenic T cells from wild-type mice and SCID mice transplanted with thymus organoids 28 days prior (n=10-15 per group). B) Proliferation and C) IFN-γ production following stimulation of splenocytes isolated from organoid recipient SCID mice. Splenocytes were isolated 28 days post-organoid transplant and labelled with CFSE proliferation dye. Cells were stimulated with CD3/28, Con A or PHA for 96h and T cell proliferation and IFN-γ levels were quantified (n=5).

To examine whether the organoid-derived T cells were functional, we stimulated splenocytes from SCID mice that had been transplanted with thymic organoids. T cells were stimulated in culture with mouse T cell activator CD3/CD28 dynabeads, 2.5ug/ml PHA, or 2.5ug/ml Con A. Organoid-derived T cells proliferated (Figure 5B) and secreted IFN-γ (Figure 5C) in response to stimulation.

### Rare T cells

The majority of mature T cells from the thymus are conventional αβ T cells, however, the thymus is also the primary site for the development of rare immune cells such as Tregs, γδ T cells, and NKT cells. We were able to detect both FOXP3^+^ Tregs (Figure 6A) and γδ T cells (Figure 6B) in mice that received thymic organoids. In addition, transplanted thymic organoids also reconstituted SCID mice with NKT cells (Figure 6C). Collectively, the thymic organoids give rise to a variety of T cells, similar to the native thymus.

**Figure 6.**
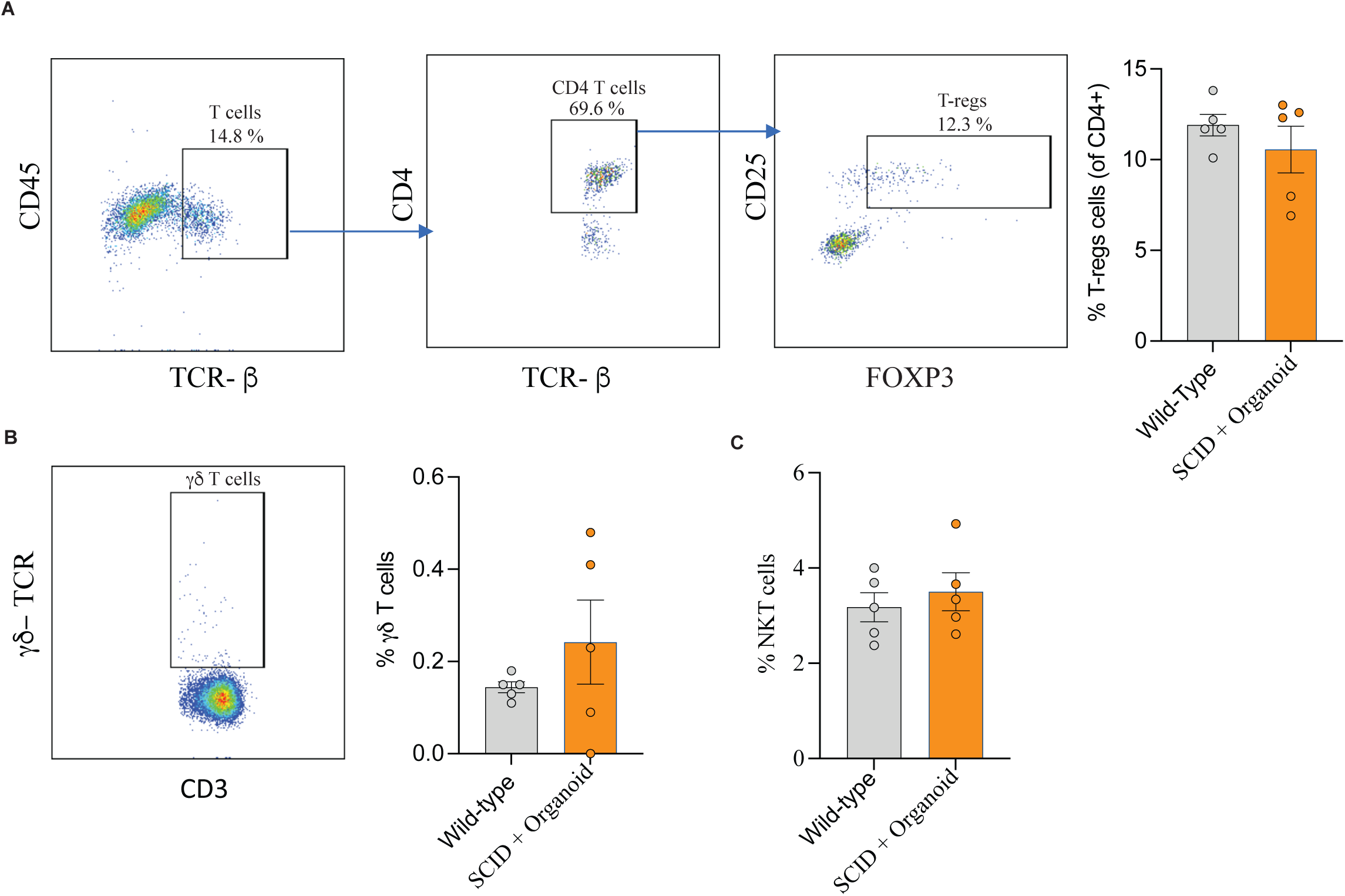
Thymus organoids gave rise to CD4^+^ FOXP3^+^ T cells, δγ T cells and NKT cells. Splenocytes from wild-type or organoid recipient SCID mice were stained for A) Tregs (CD4^+^ CD25^+^ FOXP3^+^), γδ T cells (CD3^+^εδ TCR^+^) and C) NKT cells (CD3^+^ NK1.1^+^) (n=5). Splenocytes were isolated 28 days following organoid transplantation and the T cell subsets were evaluated using flow cytometry.

### Tissue distribution and organoid staining

To examine T cell distribution in tissues, we quantified the relative expression of CD3ε transcripts compared to β-actin expression across different tissues. The levels of CD3ε in SCID mice were below the threshold of detection, whereas mice receiving thymus organoids had CD3ε detected in the spleen, liver, heart, lung, and kidneys (Supplementary Figure 3).

In addition, approximately 3 weeks following organoid transplantation, we observed a single organ-like structure forming at the site of subcutaneous transplantation (Supplementary Figure 4A). When we stained for T cells with anti-CD3, we observed more T cells in the periphery of the organoid compared to the core (Supplementary Figure 4B). In contrast, fibroblasts (Vimentin^+^) were widely distributed across the tissue (Supplementary Figure 4C). We did not see any clear medulla/cortical separation that is typically observed in the developed thymus.

### Tumor model

To examine whether thymus organoids can generate functional T cells, we employed the pmel-1 mice that produce gp-100-reactive Vβ13^+^ T cells (23). These T cells have been shown to protect mice from B16 Melanoma growth. We generated thymus organoids using thymic cells from pmel-1 mice. The pmel-1 organoids were transplanted into wild-type mice, and one week later, the mice were inoculated with B16 melanoma (Figure 7A). This delayed inoculation was necessary to enable the thymus organoids to produce a sufficient number of Vβ13^+^ T cells. Mice transplanted with pmel-1-organoids showed significant protection from tumor growth starting 10 days post-tumor inoculation (Figure 7B). This protection was associated with a higher infiltration of CD45^+^ immune cells, T cells, NK cells, and Vβ13^+^ T cells in mice that received pmel-1-organoids compared to control mice (Figure 7C, Supplementary Figure 6). We did not observe any differences in CD69 expression on the infiltrating immune cells (data not shown). CD69 is upregulated early on activated T and NK cells, and CD69 expression is transient and gets downregulated within a few hours. In contrast, the draining lymph nodes from mice transplanted with pmel-1 organoids had fewer CD45^+^ immune cells, T cells, and Vβ13 T cells compared to vehicle controls, and no difference in NK cell numbers (Figure 7D). This is likely due to increased migration of immune cells from the draining lymph nodes to the tumor. However, mice that received pmel-1-organoids also had a higher frequency of T cells, Vβ13 T cells, and NK cells expressing CD69 compared to vehicle control (Figure 7D). This indicates enhanced immune activation within the draining lymph nodes of mice transplanted with pmel-1 organoids. Collectively, the data suggest that pmel-1 organoids give rise to functional antigen-reactive Vβ13^+^ T cells that protect mice from tumor growth. Mice receiving thymus organoids also showed enhanced immune activation within the draining lymph nodes and increased tumor immune infiltration. NKT cell frequencies in the tumor and draining lymph nodes were too few to enumerate reliably.

**Figure 7.**
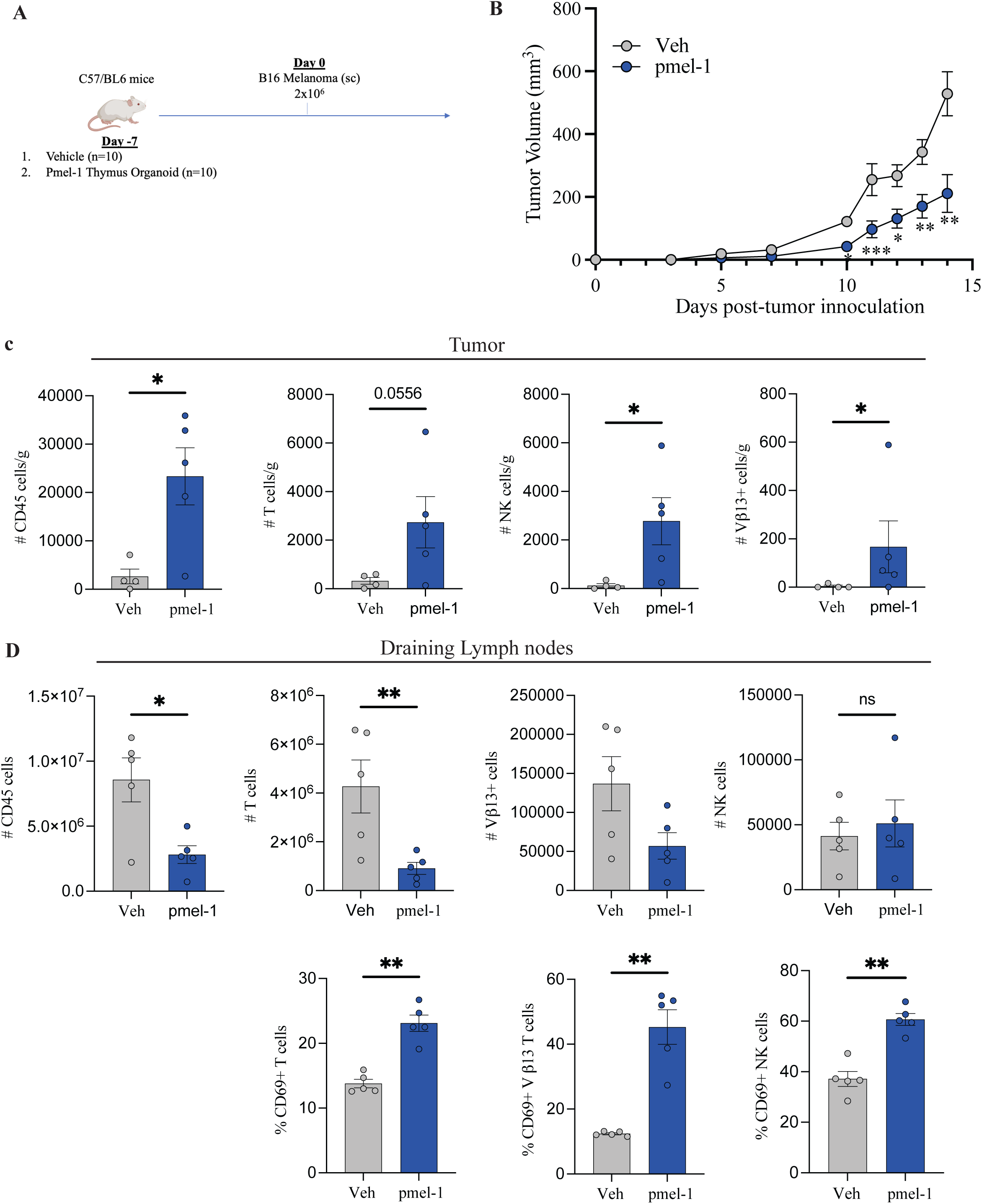
Pmel-1 thymus organoids delayed B16 melanoma growth and enhanced tumor immunity. A) Schematic subcutaneous B16 melanoma model. Wild-type C57/BL6 mice were injected subcutaneously with either vehicle or pmel-1 thymus organoid. One week later, mice were inoculated with 2×10^5^ B16 melanoma cells resuspended in Matrigel (n=10). B) Tumor volume was measured for 14 days following inoculation. C) Tumors and E) Draining lymph nodes were harvested on day 14, and flow cytometry was used to assess accumulation of immune cells (n=4-5). The frequency of CD45^+^ leukocytes, T cells (CD3^+^), NK cells (CD3^-^ NK1.1^+^), Vβ13^+^ CD3^+^ T cells, and the expression of early activation marker CD69 was examined (n=5). Total cell counts were performed on the NC3000 cell counter, and immune cell infiltration was normalized to tumor’s mass in grams.

## Discussion

Owing to progress in three-dimensional cultures, we have significantly advanced discoveries surrounding organ development and organoid formation. Organoids derived from stem or progenitor cells have become well established and are rapidly approaching the clinical testing phase (24). Despite these encouraging trends, there has been limited clinical progress in immune system-related organoids, in part due to the complex nature of crosstalk between the stromal compartment and immune cells required. Here, we have developed a rapid, repeatable, and scalable process for producing thymic micro-organoids. The thymus organoids can be injected subcutaneously and give rise to mature and functional T cells when transplanted into immunodeficient mice. The organoids persist for months after transplantation (Supplementary Figure 4).

Historically, two-dimensional TEC cultures have had limited success in generating functional T cells (8). Zuniga-Pfluckers’ group developed a widely used T cell development culture using hematopoietic stem and progenitor cells (HSPCs) with OP9 cells expressing notch ligands (7, 10). This approach was later optimized for iPSCs (25). Seet and colleagues further advanced the thymus organoid field when they developed 3D cultures using MS5 cells expressing notch ligands and HSPCs from various sources (26). They later demonstrated that iPSCs could also be differentiated into mature T cells using this system (9, 18). While these approaches have primarily sought to replace the need for thymic epithelial cells in organoids, there have also been advances in isolation and/or differentiation of thymic epithelial cells from various sources. Recent reports of 3D thymic organoids using native thymic epithelial cells or iPSC-derived epithelial cells have sought to address the gaps in 2D thymic/TEC cultures (14, 15, 27–29). Ramos *et. al* described a process of differentiating TECs from stem cell-derived thymic organoids *in vitro* (14), marking an advancement of previous studies that showed iPSC differentiation to thymic epithelial progenitors (28–30). However, maintaining these cultures requires cumbersome culture techniques and the long-term functionality of these epithelial organoids needs to be well studied. Recent publications demonstrate that TEC organoids derived from embryonic or post-natal thymus can be cultured under minimal requirement conditions and maintain their ability to support T cell development (11). Lim and colleagues also demonstrated that TEC organoids derived from adult mouse thymus could be cultured for more than 2 years, maintaining their phenotype and function (15). There is ongoing work from the same group to develop TEC organoids from adult human thymus (unpublished work).

While this manuscript was being prepared, a recent publication used a similar approach to develop a microwell-based miniaturized thymic organoid culture (13). In their body of work, they optimized thymic reaggregation cultures in coated micro-wells over 14 days in culture. Similar to our approach, the miniature organoids are well-suited for high-throughput studies (13). Our approach uses exogenous fibroblasts to expedite the formation of thymus micro-organoids, which can be cryopreserved or transplanted subcutaneously within 3 days of culture. We observed fewer variabilities compared to those acknowledged by the authors, and the shortened duration of our cultures also negates the decline in thymocytes during extended culture (13). Similar to their findings, our thymus organoids were able to give rise to αβ T cells (including NKT cells) and γδ T cells. While the authors did not investigate whether their miniature organoids can give rise to FOXP3^+^ Tregs, they reported an underrepresentation of Autoimmune regulator (Aire)^+^ mature mTECII population in their organoids (13). These cells play an important role in Treg development and tolerance. While characterizing Aire expression was beyond the scope of this manuscript, our thymus organoids gave rise to FOXP3^+^ Tregs when transplanted in mice. Furthermore, their findings were based on *in vitro* studies, whereas our data extends to *in vivo* functional validation (13). Collectively, both manuscripts highlight important advancements in the scale-up of thymic organoid production and potential use in both discovery and translational work.

Thymus organoids can be a powerful tool for enabling discoveries surrounding T cell development and have an enormous untapped therapeutic potential. For instance, artificial thymus organoids have been used to generate T cells from bone marrow organoid-derived precursors (31) and T cells with engineered chimeric antigen receptors (32). Our micro-organoids can further advance these therapeutic approaches, as they retain the capacity for TEC-mediated functions such as positive selection and central tolerance. We have demonstrated that our organoids can easily be scaled up, give rise to different T cell subsets, including δγ T cells, NKT cells, and Tregs, making this approach highly amenable to T cell therapeutics. Furthermore, the organoid-derived T cells are responsive to stimulation in culture and can give rise to antigen-reactive T cells that protect mice from B16 Melanoma. Future work will focus on optimizing the protocol for human thymic organoids, fine-tuning the selection of specific T cell subsets, and addressing the challenges associated with allogeneic T cell therapeutics.

## Supplementary Figure Legend

**Supplementary** Figure 1. Spatial distribution of thymus cells and fibroblasts three days post-seeding. Fibroblasts were labelled with Calcein AM dye and thymus cells with Eflour670 dye prior to seeding. After 72 hours, organoids were imaged on the Echo resolve microscope. Organoid diameter was ∼150 μm.

**Supplementary** Figure 2. Organoid-recipient SCID mice had circulating T cells within 2 weeks following transplant. A) Representative flow cytometry plots of circulating T cells (TCR^+^ CD45^+^) in SCID mice, and SCID mice transplanted with thymus organoids. Numbers indicate percentage frequency of T cells. B) Number of circulating T cells per volume of blood was examined 14 days following thymus organoid transplantation in SCID mice (n=3).

**Supplementary** Figure 3. Tissue distribution of T cells in SCID mice following organoid transplantation. CD3ε mRNA expression in the spleen, liver, lung, heart, and kidney of SCID mice and SCID mice transplanted with thymus organoids 28 days prior. Relative gene expression was analyzed by comparing to the validated housekeeping gene β-actin (2−ΔCT) (n=3).

**Supplementary** Figure 4. Transplanted micro-organoids form a large organ-like structure *in vivo*. A) Image of organ-like structure 8 weeks after injection of micro-organoids. The organoid was harvested and processed for immunohistochemistry analysis. B) CD3 (brown) and C) Vimentin (brown) staining on organoid tissue harvested 8 weeks following transplantation in SCID mice. Hematoxylin (purple) counterstaining was performed on both tissue stains. Black bars indicate 100 μm.

**Supplementary** Figure 5. Representative flow cytometry plots showing the gating strategy used to identify T cells, NKT cells, NK cells, Vβ13 T cells and CD69^+^ cells.

