## Supplementary figures and images for "A Rapid and Scalable Subcutaneously Administered Murine Thymus Micro-organoid for Generating Functional T cells"

### Supplementary Figures

S1

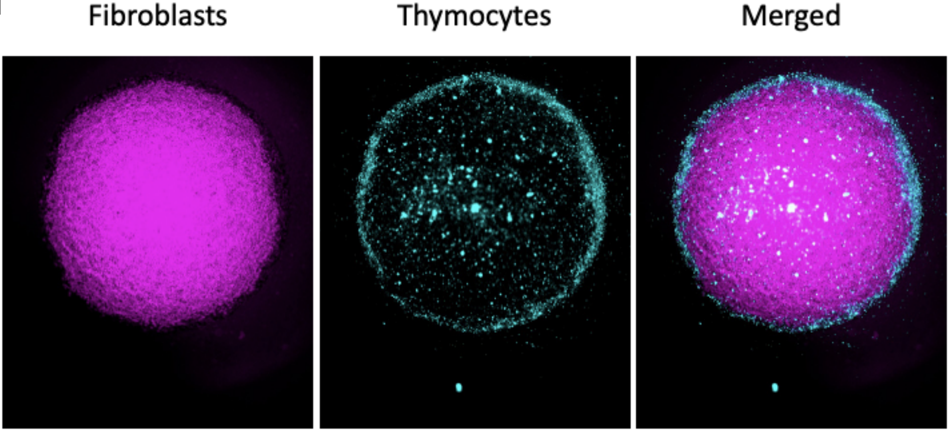

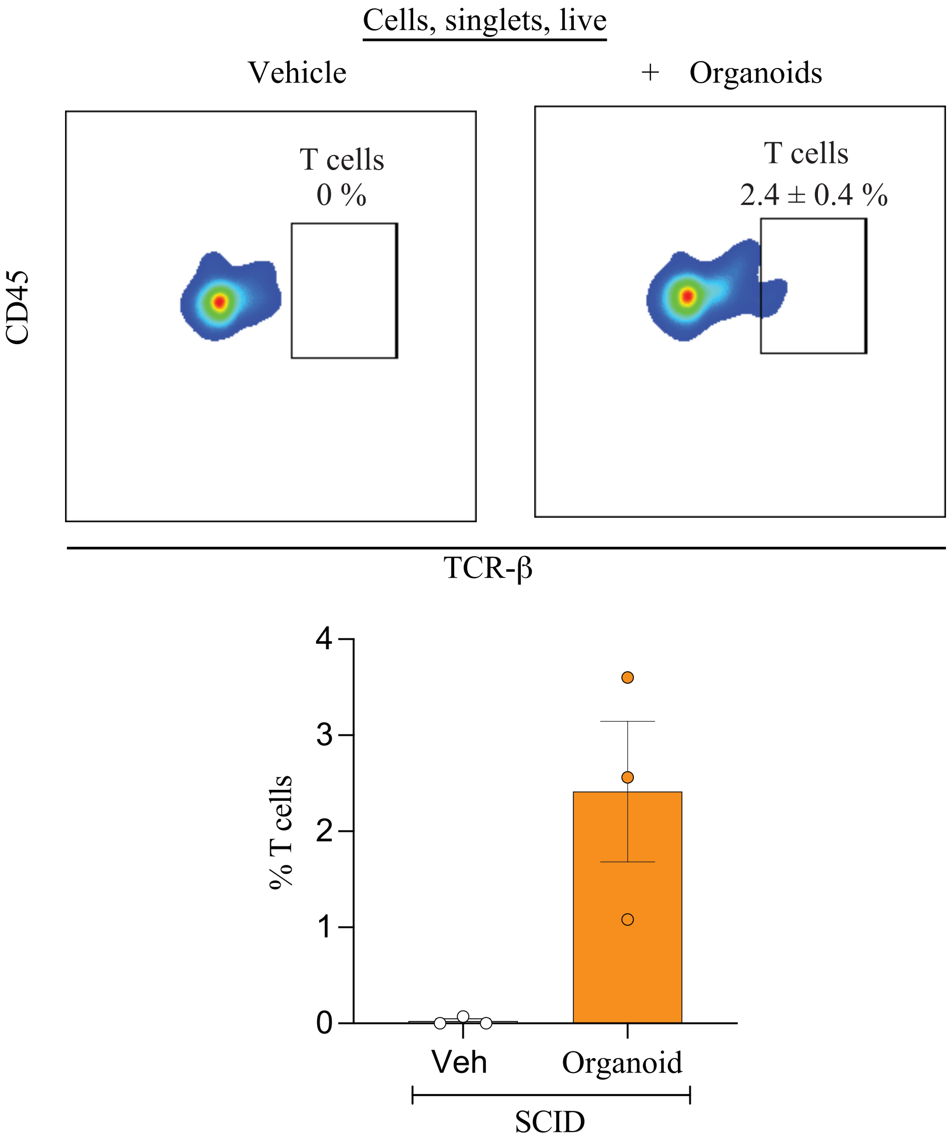

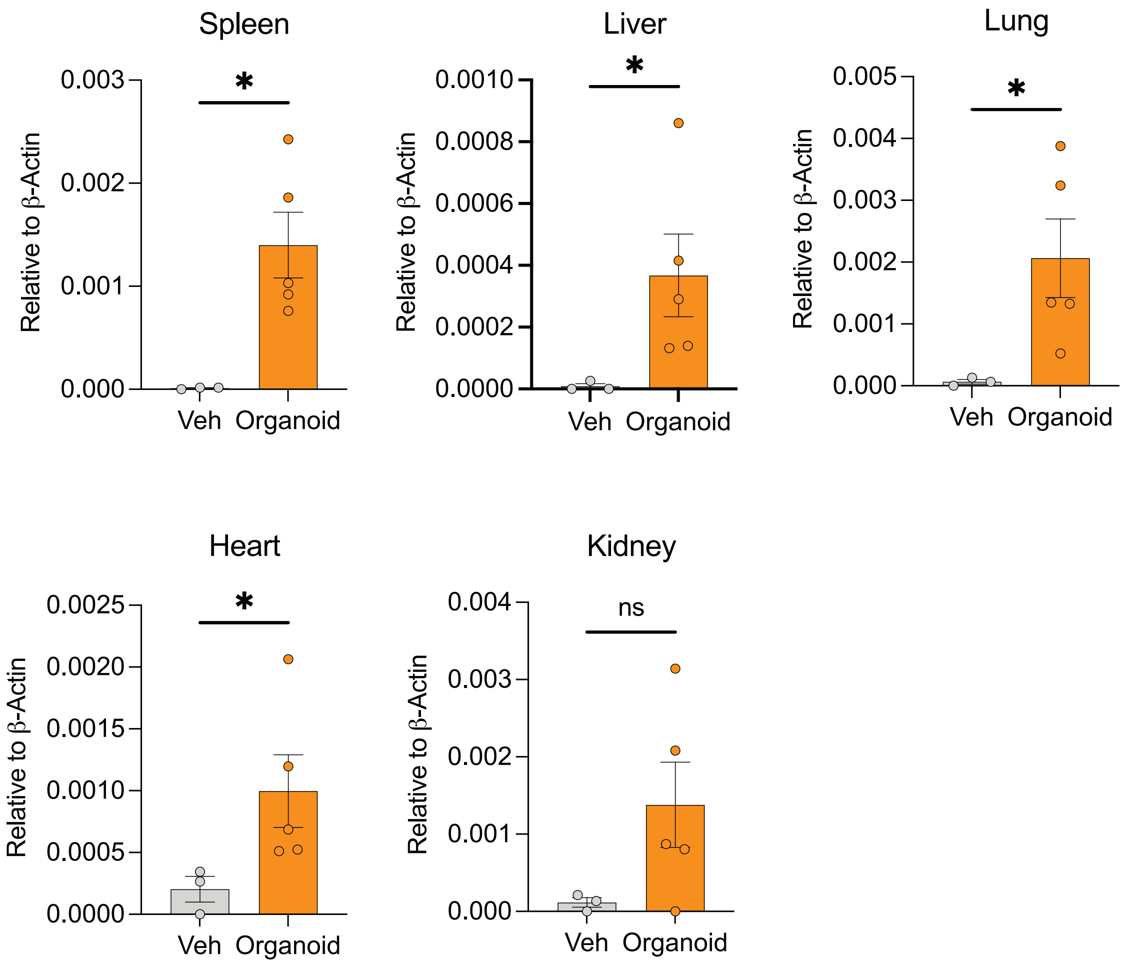

S4

A

B

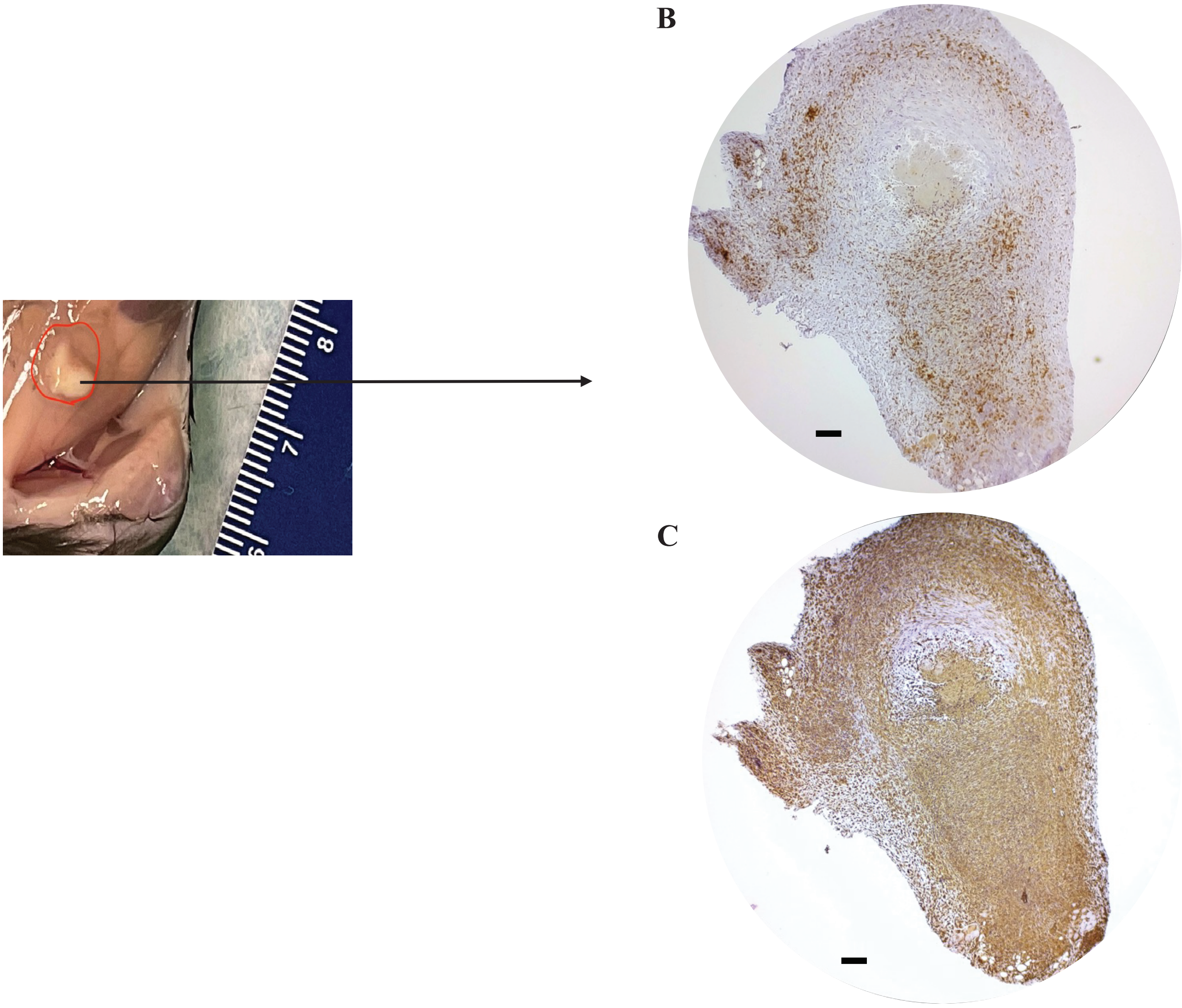

CD3

C

Vimentin

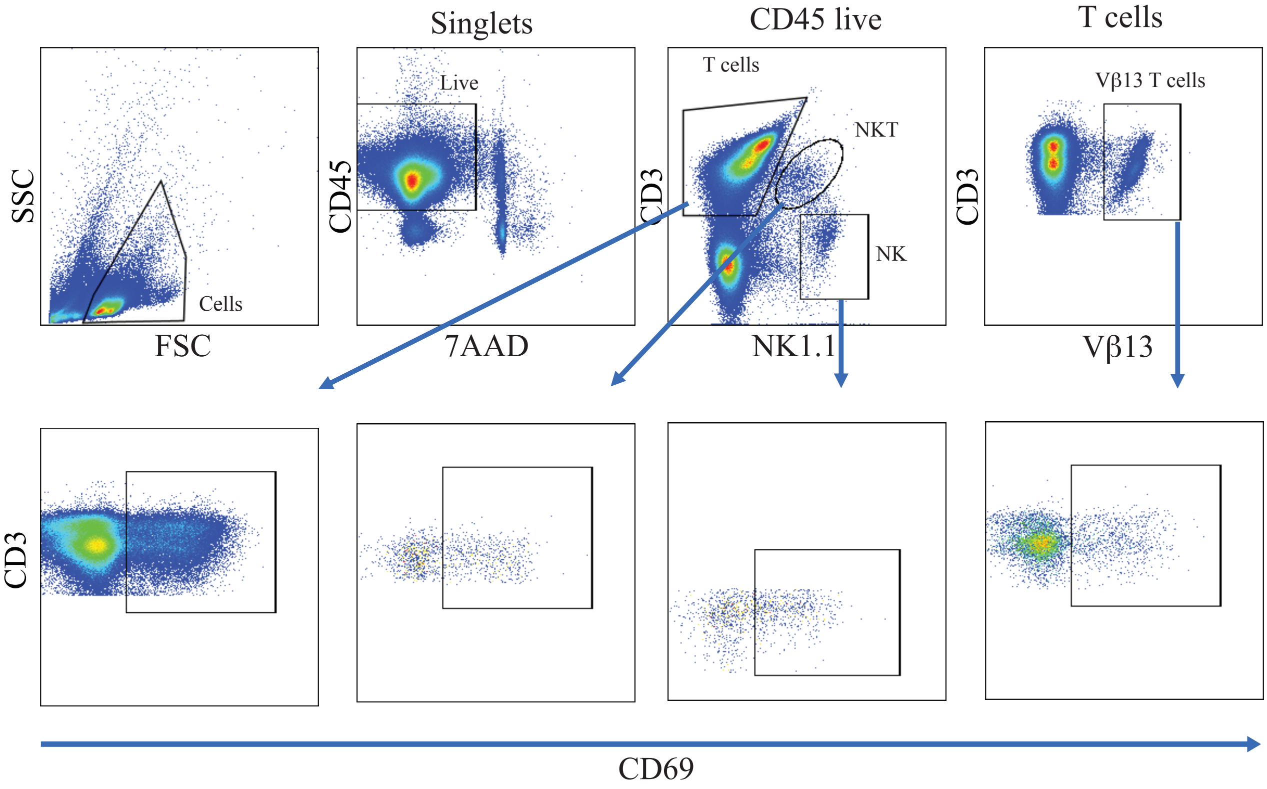
